# Metabolic early warning signals of epigenetic tipping points under chromatin modifier competition

**DOI:** 10.1101/2025.03.24.644914

**Authors:** Tomás Alarcón, Javier A. Menendez, Josep Sardanyés

## Abstract

The maintenance of epigenetic landscapes (EL) requires the precise regulation of chromatin-modifying enzymes (ChME). Competition for ChME can lead to degradation of ELs, triggering large-scale changes in the cell fate information contained in EL. Predicting impending epigenetic tipping points (ETP) by identifying early warning signals (EWS) may help to anticipate the onset of cell identity loss during aging and cancer. We have developed a general mathematical framework that incorporates different connectivity patterns generated by the 3D chromatin folding structure to analyze competition-induced ETP in large EL. This framework allows us to measure the sensitivity and robustness of ETP to the availability of metabolic cofactors and to identify potential EWS. Using a dimension reduction method, we derived coarse-grained (CG) equations for the collective observables associated with chromatin modifications. Analysis of the CG system allows the prediction of global transitions that shape the large-scale features of EL, accurately reproduce the corresponding microscopic benchmarks, and reveal the existence of tipping points under conditions of ChME competition. We applied the CG method to predict ETP under different connectivity patterns, including heterogeneous profiles such as those found in Hi-C data. Although a robustness measure for stable EL was derived from the CG dynamics in bistable regimes, sensitivity analysis revealed that metabolic cofactors have the greatest impact on EL robustness. In particular, we identified the metabolic cofactors SAM and acetyl-CoA as potential EWS for the catastrophic loss of hyperacetylated EL induced by ChME competition. The ability to predict global ETP can facilitate the discovery of predictive biomarkers and inform metabolic interventions aimed at limiting and reversing pathological cell fate decisions.

**Author summary:** Cells maintain their identity through specific patterns of gene expression controlled by structures called “epigenetic landscapes” (EL). Deterioration of these EL can occur over time as the enzymatic machineries responsible for maintaining them compete with each other. When this competition reaches a “tipping point”, cells can undergo critical shifts in cell fate and lose their original identity, leading to aging and cancer. Using mathematical models, the researchers developed a way to accurately predict these tipping points early on, based on “warning signals” derived from the abundance levels of metabolites used by chromatin-modifying enzymes. This work could lead to new methods for detecting when cells are about to lose their identity and open the door to metabolic treatments that prevent or reverse this process in aging and cancer.

## Introduction

Multicellular organisms are composed of many different cell types that perform specialized functions and are characterized by specific patterns of gene expression. Due to intrinsic stochasticity and environmental cues, heterogeneity can compromise gene expression patterns associated with specific cell identities. To prevent a cellular identity crisis, cells have evolved several regulatory mechanisms that tightly control gene expression, involving transcriptional, translational, post-translational, and epigenetic elements. In particular, maintaining stable epigenetic landscapes (EL) is essential to confer specific fates and functions to cells and requires the timely regulation of chromatin modifiers, including epigenetic enzymes and their associated metabolic cofactors. Dysregulation of chromatin-modifying enzymes and metabolic fluxes could drive the epigenetic regulatory system through “tipping points” that induce sudden and large transitions in the EL. Such critical transitions can fundamentally alter both the structure and the information contained within the EL, thereby leading to changes in cellular identity. The deterioration of the information storage in (and retrieval from) the EL may drive physiological responses such as age-related tissue dysfunction [1] or key phenotypic switches such as the epithelial-to-mesenchymal transition (EMT), which plays important physiological roles in embryonic development, tissue regeneration, and wound healing, but also occurs in pathological processes such as cancer progression and organ fibrosis [2].

Global EL transitions can be induced in response to variations in chromatin modifiers [3] or changes in chromatin structure due to dynamic rewiring of the interaction map between genome regions [4]. Competition for chromatin modifiers has been identified as a candidate for inducing large-scale global transitions, where large changes in ELs occur in response to small changes in a given regulatory state.

According to the “relocalization of chromatin modifiers hypothesis,” the accumulation of DNA damage may lead to excessive competition for dual-function epigenetic enzymes such as sirtuins, which facilitate DNA repair at DNA double-strand breaks (DSBs), but also help to epigenetically maintain specific patterns of gene expression, particularly at developmental genes [5]. The establishment of a positive feedback loop of chromatin modifications in response to DNA damage has been proposed to promote the erosion of ELs and accelerate the loss of cellular identity that drives aging [6]. Another example of global transitions in ELs governed by competition between histone modifications and transcription factor dysregulation is the EMT phenotypic switch in which epithelial cells lose cell-cell adhesion and apico-basal polarity to gain migration and invasion properties [2]. Such qualitative transitions in the ELs may indeed represent tipping points, i.e., sudden transitions that alter the global structure of an EL.

All of the above evidence highlights the emerging recognition that competition between multiple sources of epigenetic variability (such as chromatin structure, histone modifications, transcription factor binding, etc.) is a key driver of EL transitions and maintenance versus loss of epigenetic information. However, we lack a general theoretical framework to analyze the behavior of (competition-induced) tipping points in large-scale ELs, to identify the sensitivity and the robustness of such ELs to epigenetic variability such as the fluctuations in metabolic cofactors, and to potentially identify early warning signals (EWS). EWS, which typically arise when a nonlinear system approaches a tipping point, involve changes in the dynamics associated with variability, memory, return times, and flickering [7, 8]. Although EWS have been extensively studied in complex ecological and global climate systems [9–14], their use in predicting the onset of disease is only beginning to be explored as evidence accumulates that in complex diseases such as cancer, autoimmunity, cystic fibrosis, AIDS, influenza, and others, deterioration is typically abrupt rather than smooth [15–17]. Like ecosystems, molecular biosystems exhibit qualitative transitions in their response to stochastic perturbations but show little sign of the impending switch from a healthy to a diseased state until it has occurred. Therefore, to predict these critical transitions, it is crucial to identify EWS capable of capturing the proximity to tipping points.

Patterns of chromatin modifications have been studied using theoretical models in small genomic regions on the order of a few kilobases [18], and, less frequently, in larger systems [19]. These models are computational in nature, and their analysis for stability or parameter sensitivity can only be performed by extensive (and costly) numerical simulation. As a first step to remedy this situation, we address here the development of a framework that allows for analytical and feasible computational approaches. Using dimension reduction techniques, we derive coarse-grained (CG) evolution equations for the collective observables associated with specific chromatin modifications. By analyzing the CG model, both in the stochastic and mean-field regimes, our framework can predict abrupt, global transitions in the large-scale features of ELs (see Fig. 1 and Fig. S2 in the Supplementary Materials for an illustration of the analysis pipeline).

**Fig 1.**
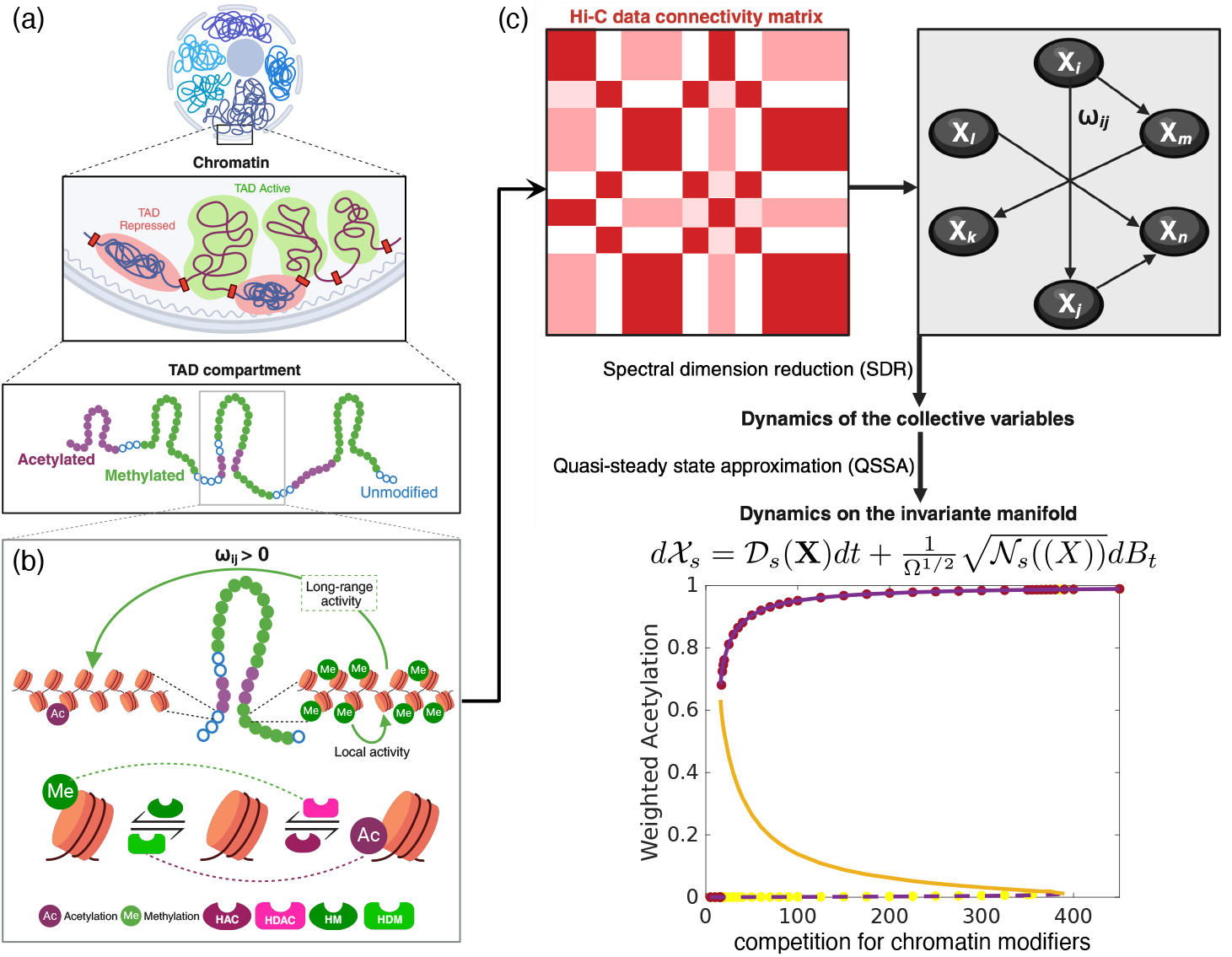
Schematic representation of the model of chromatin modifications with local and long-range activity. (a) DNA is organized in a complex, multiscale fashion, from chromosomes to sequence. Here, we focus on the scale of topologically-associating domains (TADs), which are regions with higher-than-average inner connectivity and which are loosely connected with genomic regions outside the TAD. TADs exhibit a rather large degree of variation in sizes, ranging between 100-5000 kb [20, 21]. Based on quantitative information for Hi-C maps, the resolution of such maps is of the order of 1 kilobase (kb) [22]. Since each nucleosome is wrapped with DNA of approximately 150 bases in length, each region in our model contains between 5 and 10 nucleosomes. Since each nucleosome contains two H3 histones, this implies that the number of H3K27 sites available for modification, *S*_0_, is of the order of 10-20. (b) Negative and positive feedback on chromatin modifiers by epigenetic modifications play a crucial role in generating and maintaining stable epigenetic landscapes. The feedbacks that we are incorporating in our model are shown in the lower panel of (b) (adapted from Dodd et al. [23], see also [2]). As illustrated in (b), such feedbacks can be either *cis* (local, within a region) or *trans* (non-local, between regions). We assume that the network of physical contacts between regions (represented here by the dashed line) provides the backbone for such feedbacks. (c) Schematic representation of the modeling pipeline where the microscopic system is reduced using first a spectral dimension reduction method and then a quasi-steady state approximation. The result is a low-dimensional system of stochastic differential equations for a set of collective variables, whose critical behavior accurately reproduces that of the full microscopic system. Full details are provided in Materials & Methods and in the Supplementary Materials, see also Fig. S2.

In particular, we show that the predictions of the CG model robustly and accurately reproduce those of the corresponding microscopic benchmark. Specifically, we demonstrate the existence of tipping points under conditions of competition for chromatin modifiers. In multi-stable regimes, the asymptotic analysis of the (stochastic) low-dimensional, coarse-grained dynamics uniquely allows us to derive a novel robustness measure for stable ELs. Our framework integrates several biological variables including competition for chromatin modifiers, metabolic cofactors, and the connectivity patterns generated by the 3D folding structure of chromatin on the robustness of the EL. Because our framework can be treated analytically, we can gain a deeper understanding of how these three components interact not only in the generation of robust EL, but also in EL erosion leading to loss of epigenetic information. By performing a sensitivity analysis of the robustness measure, we extend our analysis to determine the model parameters with the greatest impact on the loss of robustness for specific ELs. Such an analysis allows us to identify that the availability of two metabolic cofactors for chromatin-modifying reactions, SAM and Acetyl-CoA, strongly influences the robustness of specific ELs and thus correlates closely with EWS for predicting the proximity of EL tipping points. Our methodology may help to guide metabolic interventions to prevent or even reverse undesired global EL transitions that underlie the collapses of cellular identity in aging and cancer.

## Materials and methods

We provide a brief discussion of the methodology used to model the epigenetic modifications within a genomic region (see Fig. 1), the coarse-graining procedure that renders the model tractable, and, finally, the tools for analysis of robustness and parameter sensitivity. Full details and derivations are provided in the Supplementary Materials.

### Mathematical model of chromatin modifications

We consider a simplified model that solely takes into account modifications of H3K27, specifically acetylation and methylation [18, 23]. We formulate a stochastic dynamical system for the number of acetylated and methylated residues within a specific region. We number such regions according to *i* = 1, …, *N*_*C*_, where *N*_*C*_ is the total number of regions considered in our simulations (see Fig. 1(a)). Chromatin modifications involve the addition and removal of covalent modifications and, therefore, are mediated by enzymes. Thus, at each site *i* = 1, …, *N*_*C*_, we consider the Michaelis-Menten (MM) model of enzyme kinetics to model each of the chromatin modifications (i.e., addition/removal of methylation, addition/removal of acetylation) [2, 24–26]. The MM model assumes that the substrate and the enzyme form a complex that then relaxes into a product (modified chromatin) and returns the enzyme. Specifically, the addition of (tri)methylation and acetylation marks to H3K27 is modeled by:

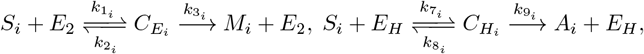

respectively. In this case, the substrates are unmodified H3K27 residues (*S*_*i*_). Methylation is mediated by EZH2, the enzymatically active member of the polycomb repressor complex, *E*_2_, while its acetylation is regulated by a histone acetyl transferase (HAT), *E*_*H*_. The complexes 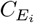 and 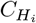, which quantify the amount of chromatin-bound enzyme, relax onto *M*_*i*_ and *A*_*i*_, which refer to the methylated and acetylated residues within bin *i*, respectively.

Demethylation and deacetylation are modeled using the same principles:

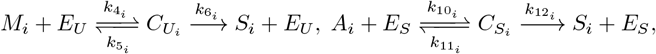

i.e., histone demethylase UTX, *E*_*U*_ bind to its substrate, *M*_*i*_, and relaxes onto its unmodified form, *S*_*i*_. The same process involving histone deacetylase (NURD or members of the SIRT family), *E*_*S*_, and substrate *A*_*i*_ leads to the removal of acetylation marks.

Furthermore, as it has been described in previous studies [19, 23, 27–29] the rate at which modifications occur in region *i* is regulated by the landscape of epigenetic marks, both locally (*cis*) and non-locally (*trans*) (see Fig. 1(a) and (b)) [18, 21]. We consider that the map of contacts provided by Hi-C datasets also provides a physical backbone for the biochemical interactions. In other words, we will assume that the modification rates at region *i* are regulated by the marks in *j* if *w*_*ij*_ *>* 0, i.e., if both regions are in physical contact. The quantities *w*_*ij*_ are the components of the matrix W = (*w*_*ij*_) and quantify the frequency with which regions *i* and *j* are in contact. We assume that such feedbacks affect the ability of chromatin modifications to facilitate or hinder the binding of the modifiers to their substrates. In general, feedback are such that acetylation (methylation) marks increase the rate at which further acetylation (methylation) occurs and inhibit the rate of addition of methylation (acetylation) marks. Specifically, feedbacks are incorporated in our model through the rates of binding and unbinding of the epigenetic enzymes to their substrates (see Supplementary Materials, Section I, for details).

The matrix of connectivity W plays a central role in the model. It reflects the 3D folding structure of chromatin and provides a scaffold for long-range propagation of epigenetic modifications [30]. To illustrate the breadth of applicability of our method and also the effects of considering different structures, we will consider three patterns of connectivity, namely, all-to-all, first-neighbors, and heterogeneous. The all-to-all connectivity corresponds to a pattern of interactions where two sites are, on average, equally likely to be connected regardless of the distance between them. The intensity of the interaction is also (on average) distance-independent. First-neighbors interactions refer to the case where neighbor sites (along the chromatin chain) have strong interactions, whereas sites beyond first neighbors are very weakly connected.

Heterogeneous connectivity is associated with cases where some subregions are very tightly connected, whereas between-subregion interactions are very weak. Hi-C maps belong to the heterogeneous class [21, 31]. For the current discussion, we will consider that W is a constant input to the model and analyze the effect of considering different architectures. Details on how we generate the W matrices are provided in Supplementary Materials, Section II.

Finally, we consider the problem of global transitions (tipping points) induced by competition for chromatin modifiers in ELs. Such situations have been studied, for example, in the context of aging [5], where sirtuins (that have HDAC activity) are recruited to DNA-damaged sites and the epithelial-to-mesenchymal transition [2]. For concreteness, we will focus on the case in which there is competition by sirtuins. We consider a simple, binding-unbinding model of sequestration of SIRT at double-strand breaks (DSBs), where a fixed amount of DSBs, *D*, captures SIRT, i.e.,

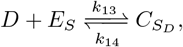

where 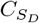 is the amount of SIRT captured by DSBs. For simplicity, we will assume that this reaction is very fast and we can consider that 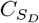 is in quasi-equilibrium.

Assuming the quasi-steady state approximation (see Supplementary Materials, Section I, for details), the mean-field equations of the microscopic model are

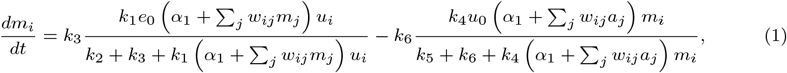

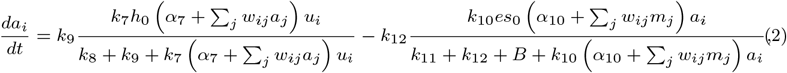

where *i* = 1, …, *N*_*C*_. Here, *m*_*i*_ and *a*_*i*_ are the levels of methylation and acetylation in bin *i, B* is the QSS rate of sequestration of SIRT by DNA-damaged sites, which is proportional to the amount of DSBs, *D* (see Supplementary Materials, Section I, for details). The stochastic formulation of the microscopic model is given in full detail in Supplementary Materials, Section I.

### Coarse-graining of the microscopic dynamics by dimension reduction

The pipeline for dimension reduction of our stochastic model is shown in SI Fig. 1. For clarity, we split its description into two parts (explained in detail in the Supplementary Materials, Section III).

The first step is based upon a dimension reduction method that reduces high-dimensional dynamical systems (such as Eqs. 1–2) to a low-dimensional version, which can then be used to describe the collective dynamics of the original system. The dimension reduction method is based on spectral graph theory, particularly on the dominant eigenvalue and eigenvector of the matrix that defines the connectivity pattern of the system [32, 33].

The second step involves the quasi-steady-state approximation (QSSA), [26, 34], also known as the stochastic averaging principle, which provides a practical way to reduce the number of degrees of freedom in the system. The key parameter that determines the form of the reduced model is the ratio of the timescale for fast variables to the timescale for the evolution of the slow variables. In our case, we apply the QSSA to the equations for the collective dynamics (coarse-grained system) derived from the spectral dimension reduction.

### Approximate spectral dimension reduction (ASDR)

Spectral dimension reduction (SDR) is a procedure that was introduced in [35] and further developed in [32, 33] as a method to coarse-grain the dynamics of deterministic, high-dimensional dynamical systems running on complex networks. Let X = (*X*_*ir*_) where *i* = 1, …, *N*_*C*_ and *r* = 1, …, *N*_*S*_. *N*_*C*_ and *N*_*S*_ are the number of nodes in the network and the number of dynamical variables, respectively, so that the dimension of the dynamical system, *D*, is *D* = *N*_*S*_ × *N*_*C*_. The (deterministic) SDR method [32] consists of considering a vector of wei ghts u = {*u*_*i*_}_*i*=1,…,*NC*_ and a set of collective (or averaged) variables 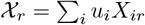 (see SI Fig. 1). For *N*_*S*_ = 1, [32] considered the set of ODEs 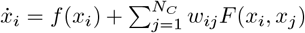. The SDR method posits that such a dynamical system on a homogeneous network can be reduced to the dynamics of the corresponding collective variable for an appropriate choice of the weights *u*_*i*_. To do so, the method proceeds to make a Taylor expansion of the functions *f* and *F* around χ and (Φ_1_χ, Φ_2_χ), respectively. It can be shown that, to the lowest order, the dynamics of the collective variable χ is given by 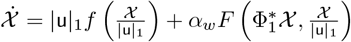 (see also Supplementary Materials, Section III). The method proposed in [32] allows us to determine in a self-consistent way the quantities u, *α*_*w*_, and 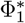, in terms of the spectral properties of the matrix W. In this work, we have extended the SDR method to a Markov-jump process (MJP). As a result, we obtain a set of stochastic differential equations (SDEs) for the collective variables (with *N*_*S*_ ≥1), thereby reducing the dimension of the problem from *D* = *N*_*S*_ × *N*_*C*_ to *D*^′^ = *N*_*S*_ ≪ *D* (see Supplementary Materials, Fig. S2). The detailed derivation of the ADSR is provided in the Supplementary Materials.

### Stochastic quasi-steady state approximation

Further dimension reduction can be achieved through the separation of time scales. Following the method introduced by Ball et al. [36] [see also [24–26]], we take advantage of the fact that some of the chemical species in the system are much more abundant than others. The associated reaction rates can vary over several orders of magnitude, inducing separation of time scales within the system so that several species evolve in much shorter time scales (fast variables) than the remaining ones (slow variables). As we show in the Supplementary Materials, Section IV, the system of SDEs for the collective variables exhibits slow-fast behavior: 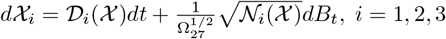, for the slow variables, and 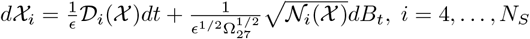, for the fast ones. The expressions of the drift and noise terms (𝒟 _*i*_ and 𝒩_*i*_, respectively) are derived in the Supplementary Materials. In order to proceed further, we use the corresponding forward Kolmogorov (Fokker-Planck) equation, which dictates the evolution of the probability density function (PDF), 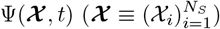 [37]:

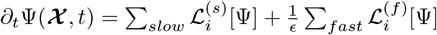, where the operators 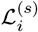 and 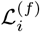 are given by 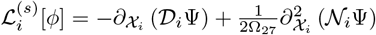 and 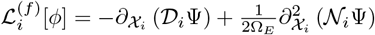, respectively. We now expand Ψ = Ψ_0_ + *ϵ*Ψ_1_ + 𝒪 (*ϵ*^2^) [34]. In the Supplementary Materials, Section IV, we show that the lowest order approximation can be written as 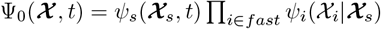 where 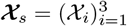 is the set of slow observables. In the Supplementary Materials, Section IV, we derive in detail the conditional (fast) PDFs *ψ*_*i*_, and, by considering the appropriate solvability conditions, we also show that, at the lowest order, *ψ*_*s*_ satisfies the *averaged* forward Kolmogorov equation

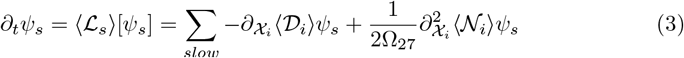

where ⟨𝒟_*i*_⟩ = ⎰_**D**_ 𝒟_*i*_*(Π*_*fast*_ *ψ*_*j*_(χ_*j*_) *d*χ_*s*_ and ⟨𝒩_*i*_⟩ = ⎰_**D**_ 𝒩_*i*_*(Π* _*fast*_ *ψ*_*j*_(χ_*j*_) *d*χ_*s*_. The rsesulting QSSA dynamics occurs in dimension 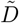 equal to that of the invariant manifold of the slow-fast deterministic dynamical system (see SI Fig. 2) with 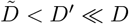. A detailed derivation of the stochastic QSSA for the ADSR equations is given in Supplementary Materials, Sections III and IV.

**Fig 2.**
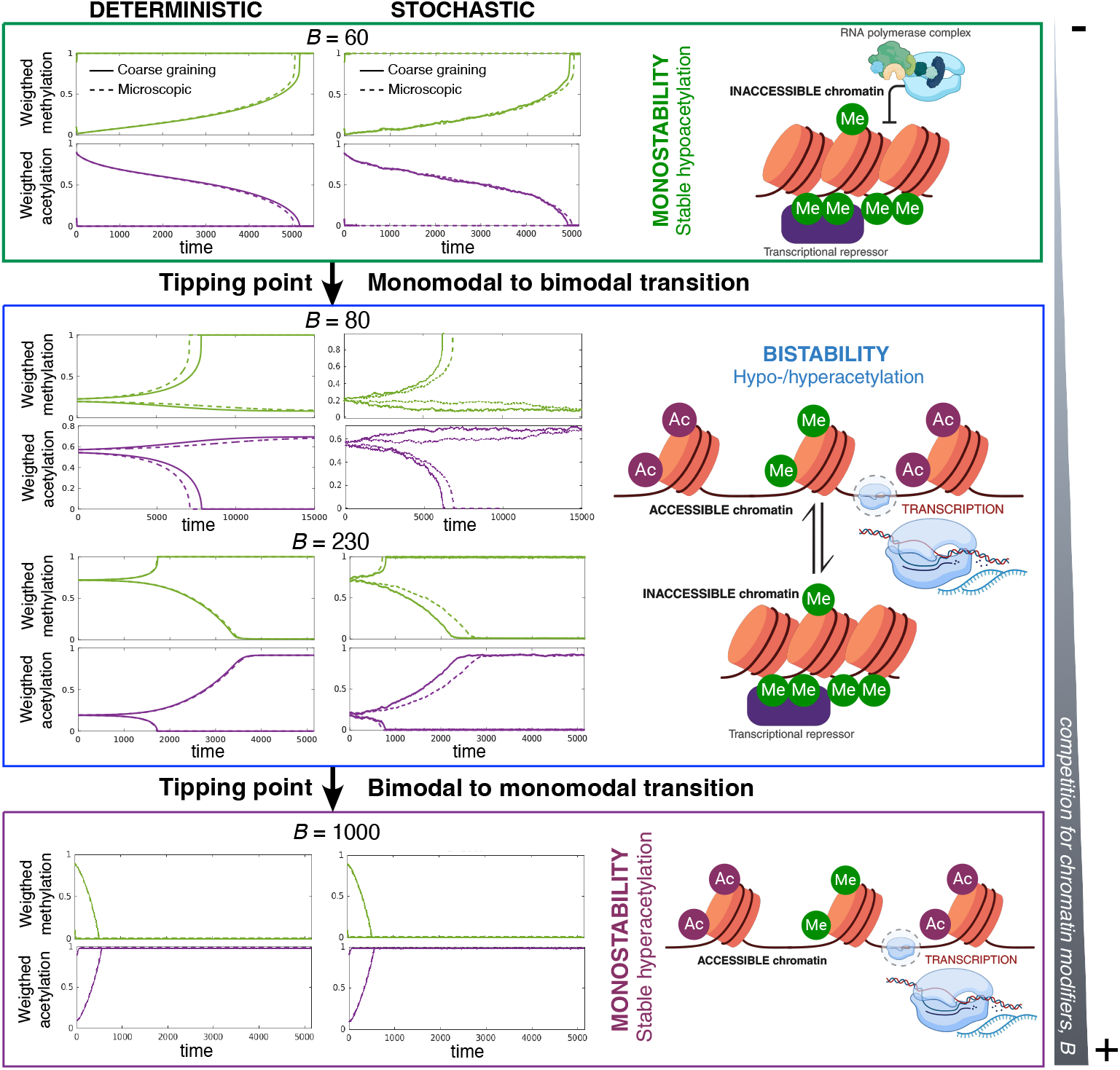
Predictions of the coarse-grained (CG) dynamics match those of the microscopic (Micro.) system, specifically regarding tipping points. Deterministic (first column) and stochastic (second column) simulations increasing the competition for chromatin modifiers, *B*. Predictions of the CG system (solid lines) predict with a high degree of accuracy the critical behavior (tipping points) of its microscopic counterpart (dashed lines). Panels display individual realizations using the all-to-all connectivity matrix.

### Robustness and parameter sensitivity analysis: A WKB approach

The analysis of tipping points and their associated early warning signals requires means to study the sensitivity of the system to changes in parameter values, specifically regarding the existence of bifurcations or phase transitions [38]. It is not uncommon for such efforts to be seriously hindered in complex, high-dimensional systems. To complete our analysis, one must obtain an analytical approximation of the solution of low-dimensional FPE. Eq. (3). To proceed further, we will assume that the system has reached a steady state, so that *ψ*_*s*_ satisfies ⟨ℒ_*s*_⟩[*ψ*_*s*_] = 0. Furthermore, since we are also assuming that Ω_27_ ≫ 1, we make a WKB Ansatz for *ψ*_*s*_, so that

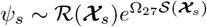 [39–43]. WKB as ymptotics prescribe that bot h ℛ and 𝒮 be expanded in powers of 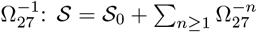 and 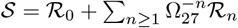. To the lowest order 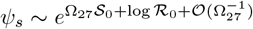, which implies that the shape of *ψ*_*s*_ strongly dominated by 𝒮_0_. According to the WKB methodology, 𝒮_0_ satisfies a Hamilton-Jacobi equation [43] (see also the Supplementary Materials): 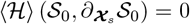, where 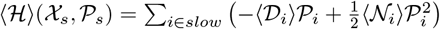, so that 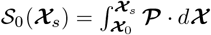, where now χ and 𝒫 are the solutions of the Hamilton equations 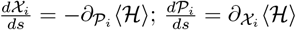, with initial conditions χ_*s*_(*s* = 0) = χ_0_ and 𝒫 _*s*_(*s* = 0) = 𝒫_0_. Because of this mathematical structure, **𝒮**_0_ is referred to as the *action* and plays a role similar to that of a quasi-potential [38].

### Robustness in bistable systems

We will now focus our analysis on regimes where the system exhibits bistability and study the robustness of the stable steady-states (SSSs). Within the context of the WKB approximation, we will analyze the resilience of SSSs to changes in parameter values. We introduce an approach to perform sensitivity analysis without the need for extensive parameter sweeps, which are computationally costly and often impractical. We will denote by 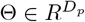 the vector whose components are the kinetic parameters of our model. *D*_*p*_ is the number of such parameters.

Specifically, we will tackle the robustness of hypoacetylated states. Let **χ**_0_ be the coordinates of the unstable saddle point and **χ**_±_, the coordinates of the stable nodes. In this context, we use as a proxy for the robustness of **χ**_±_ the quantity 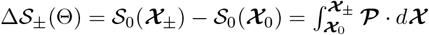. This quantity captures the features of the quasi-potential landscape associated with the FPE, Eq. (3). Specifically, it quantifies the height of the quasi-potential barrier the system needs to overcome to undergo a (noise-induced) transition from **χ**_±_ to **χ**_∓_. Δ𝒮_±_ thus provides a direct measure of the robustness of the corresponding equilibrium to intrinsic noise, and therefore, a measure of canalization of the associated epigenetic landscapes. It is therefore central to our analysis to ascertain what system parameters are those to which Δ𝒮_±_(Θ) exhibits a higher degree of sensitivity. The natural measure of sensitivity is their gradient in parameter space:

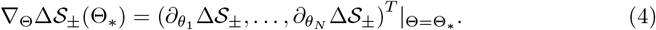

The sign and magnitude of each component of this vector provide the necessary information regarding which parameters increase the robustness of the EL and which ones decrease it. The details of the computation of this quantity are given in the Supplementary Materials.

## Results

### Coarse-grained equations

The microscopic model has been coarse-grained into the following two-dimensional system of stochastic differential equations (see the Supplementary Materials, Section III, and Fig. S2 for details):

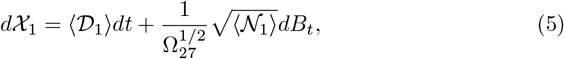

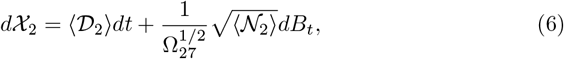

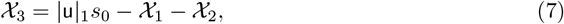

where

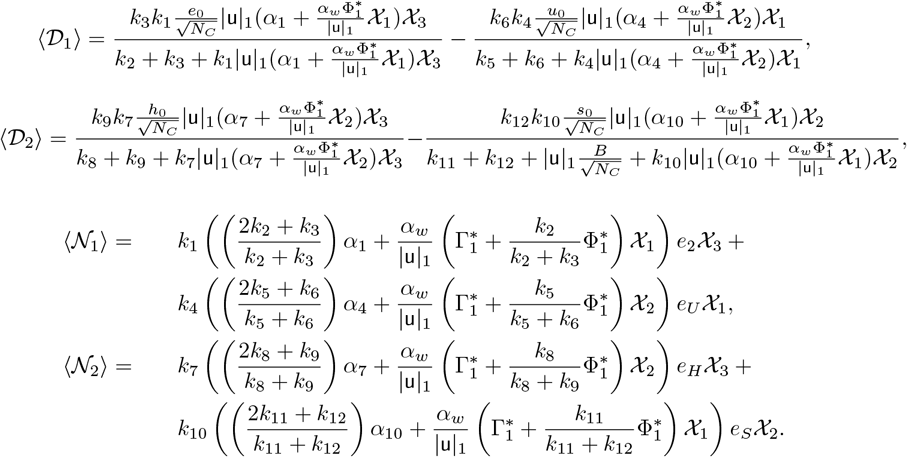

The variables χ_1_, χ_2_, and χ_3_ are the observables (weighted variables) corresponding to acetylated, methylated, and unmodified chromatin, respectively. A full description of the derivation of Eqs. (5)–(7) is given in the Supplementary Materials. Note that the form of the mathematical expressions of ⟨𝒟_1_⟩ and ⟨𝒟_2_⟩ match those of Eqs. (1)–(2), i.e., they are the addition of two Michaelis-Menten-like functions, one for addition and one for removal. The parameters appearing in the CG expressions are derived for the microscopic model in a systematic way described in detail in the Supplementary Materials, Section III, particularly the parameter 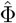, which depends on both the topology of the connectivity pattern and the strength of the contacts. It is also noteworthy the dependence of the right-hand side of Eq. (6) on the parameter *B*. As described in detail in the Supplementary Materials, Section I, *B* is a proxy for the amount of sirtuins sequestered by substrates different from chromatin, e.g., DNA damage [5]. It, therefore, quantifies the competition between chromatin and the other substrates of histone deacetylases (HDACs), specifically, sirtuins.

Figure 2 shows numerical simulations comparing the predictions of the CG system with those of the microscopic dynamics with the all-to-all connectivity matrix. In these simulations, we have used increasingly larger values of *B*, corresponding to elevating the levels of competition for the HDACs. The predictions of the CG dynamics match those of the microscopic system, both in the mean-field, deterministic scenario and in the stochastic regime. A thorough comparison between the CG and the microscopic benchmark is provided in the Supplementary Materials, Figs. S3-S6.

### Competition for chromatin modifiers triggers tipping points

The coarse-grained (CG) version of the epigenetic landscape model, Eqs. (5)–(7), allows us to analyze the system regarding the existence of transitions that alter the global structure of the landscape. Specifically, by analyzing the deterministic limit of the CG system, we address how competition for chromatin modifiers affects (HDACs in particular) the existence and stability of hypo- and hyper-acetylated landscapes. Such landscapes correspond to the steady-state observables **χ**_−_ and **χ**_+_, respectively. Our model predicts that as the level of competition for HDACs, *B*, increases, the landscape goes through several transitions (bifurcations). For intermediate values of *B*, the system is bistable, so that the hypo- and hyper-acetylated states coexist. As *B* decreases, the system approaches a tipping point (saddle-node bifurcation) where the hyperacetylated landscape ceases to exist (see Fig. 3). Conversely, if *B* increases, a second tipping point exists where the hypoacetylated state vanishes through a saddle-node bifurcation (see Fig. 2). It is noteworthy that the existence of the tipping points and the critical level of competition for HDAC (*B*) at which they occur predicted by the CG system are in excellent agreement with the results of the microscopic dynamics (see Fig. 2 and the Supplementary Materials for further information).

**Fig 3.**
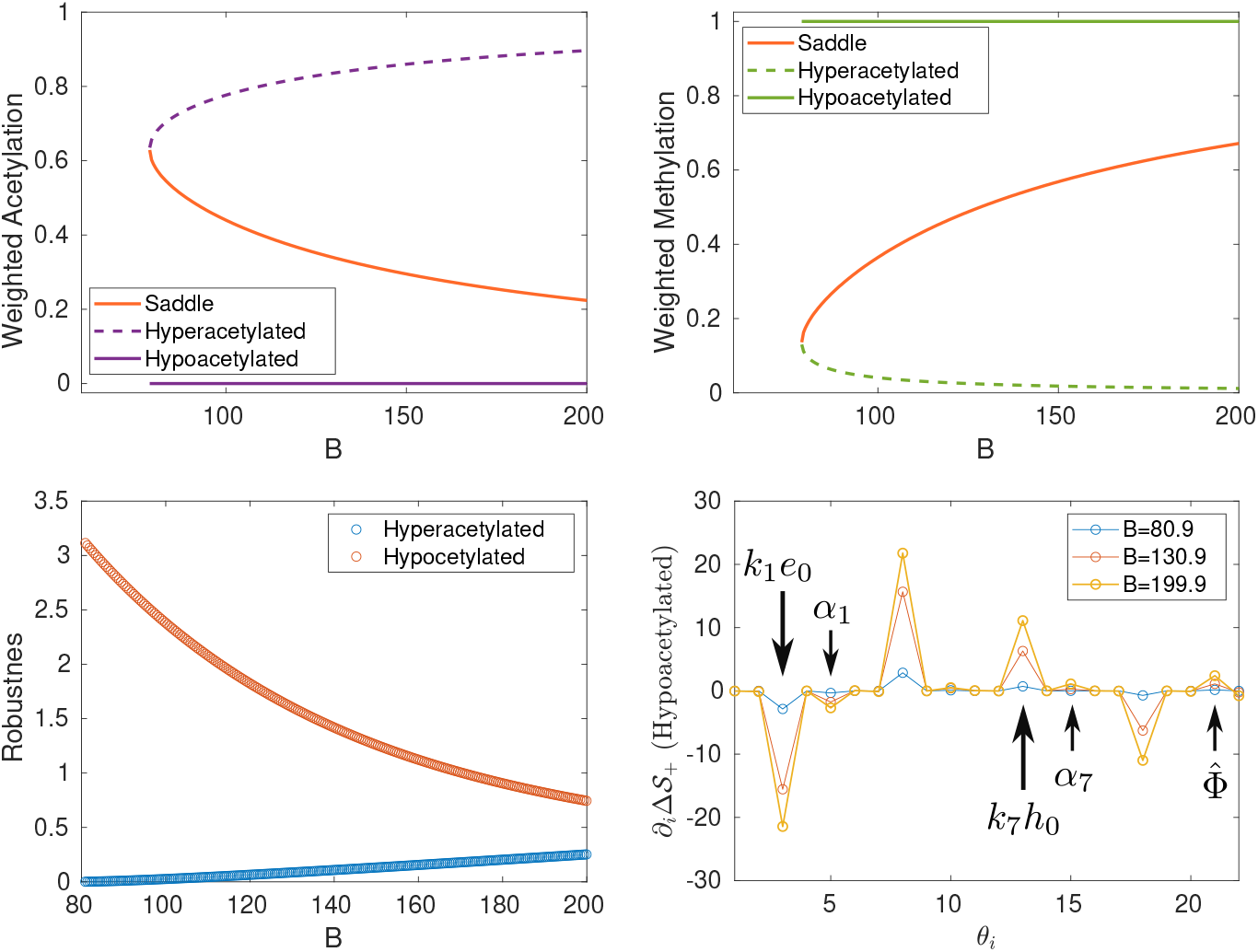
Mean-field limit and stochastic coarse-grained (CG) approximations: tipping points and sensitivity analysis. The deterministic limit of the CG approximation predicts that competition for histone deacetylases (quantified by the parameter *B*) induces tipping points in the stable ELs. The stochastic analysis allows us to quantify the robustness of the stable ELs and, more interestingly, the sensitivity of the robustness when the model parameters are changed. This sensitivity analysis allows us to determine which parameters are more effective at increasing or decreasing the robustness of the associated ELs. Legend: 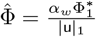. The quantities Δ***𝒮***_∓_ defined in the main text are used as proxies for the robustness of hyper- and hypo-methylated states, respectively. This analysis has been done in the case of all-to-all connectivity.

### Robustness and sensitivity analysis

As discussed in the Materials & Methods, the quantities Δ**𝒮**_∓_ are proxies of the robustness of hyper- and hypo-methylated states, respectively, as long as we restrict our study of the epigenetic landscape (EL) to its bistable regime. Taking advantage of the fact that the stochastic low-dimensional representation, we use WKB asymptotics to compute the so-called action between the unstable (saddle) equilibrium and the stable (nodes) equilibria, corresponding to the hypo- and hyper-acetylated landscapes. By capturing the features of the quasi-potential landscape associated with the stochastic dynamics of the CG system, these quantities quantify the height of the quasi-potential barrier that the system has to overcome to leave the basin of attraction of one stable EL and move into the other.

Figure 3 shows that, as the system approaches the tipping point where the hyperacetylated state ceases to exist, its robustness tends to zero. By contrast, in the vicinity of such a transition, the robustness of the hypoacetylated landscape approaches its maximum value.

This notion of robustness allows us to explore the sensitivity of ELs to changes in specific parameters. In a nutshell (more details are given in the Materials & Methods), we can use Eq. (4), to identify the directions of the space of parameters in which the gradients of Δ**𝒮**_∓_ are steeper. Such directions correspond to the parameters whose change produces larger variations in robustness. The result of this analysis is shown in Fig. 3. Not surprisingly, the steeper directions correspond to *k*_1_*e*_0_, *k*_4_*u*_0_, *k*_7_*h*_0_, and *k*_10_*s*_0_, i.e., to those parameters that are proportional to the total abundance of each of the chromatin modifiers (HMs, HDMs, HACs, and HDACs respectively).

Beyond this general observation, by looking specifically at the sensitivity to parameters related to the activity of HMs and HACs, one uncovers an interesting pattern. Besides the elevated sensitivity to *k*_1_*e*_0_ and *k*_7_*h*_0_, Fig. 3 shows that *α*_1_ and *α*_7_ (which correspond to the basal, unregulated, HM and HAC activity, respectively) are second in their effect on the robustness of hypoacetylated landscape. It is noteworthy that the robustness is also sensitive to the structure of the connectivity matrix W (connectivity pattern, intensity of interactions, etc.), as revealed by the effect of 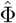 on the robustness.

### The 3D folding structure of chromatin affects the robustness of hyper- and hypo-acetylated epigenetic landscapes

Our sensitivity analysis (see previous section) reveals an interesting property of the EL-generating system, namely, that the 3D folding structure of chromatin influences robustness. Specifically, the CG sensitivity analysis shows that variations of the parameter 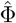, which is determined by both the structural features of the connectivity pattern and the strength of the interactions between sites, have a strong effect on the robustness of hypo- and hyper-acetylated ELs (see Fig. 3). To further explore this issue, we consider different connectivity patterns. For concreteness, we performed this analysis in the context of tipping induced by competition for sirtuins. Results of the bifurcation analysis for the mean-field limits (both microscopic and CG) are shown in Fig. 4(a-c). It is noteworthy that both the steady-state values and the critical behavior (tipping points) of the microscopic system are accurately captured by the CG system, regardless of the connectivity structure.

**Fig 4.**
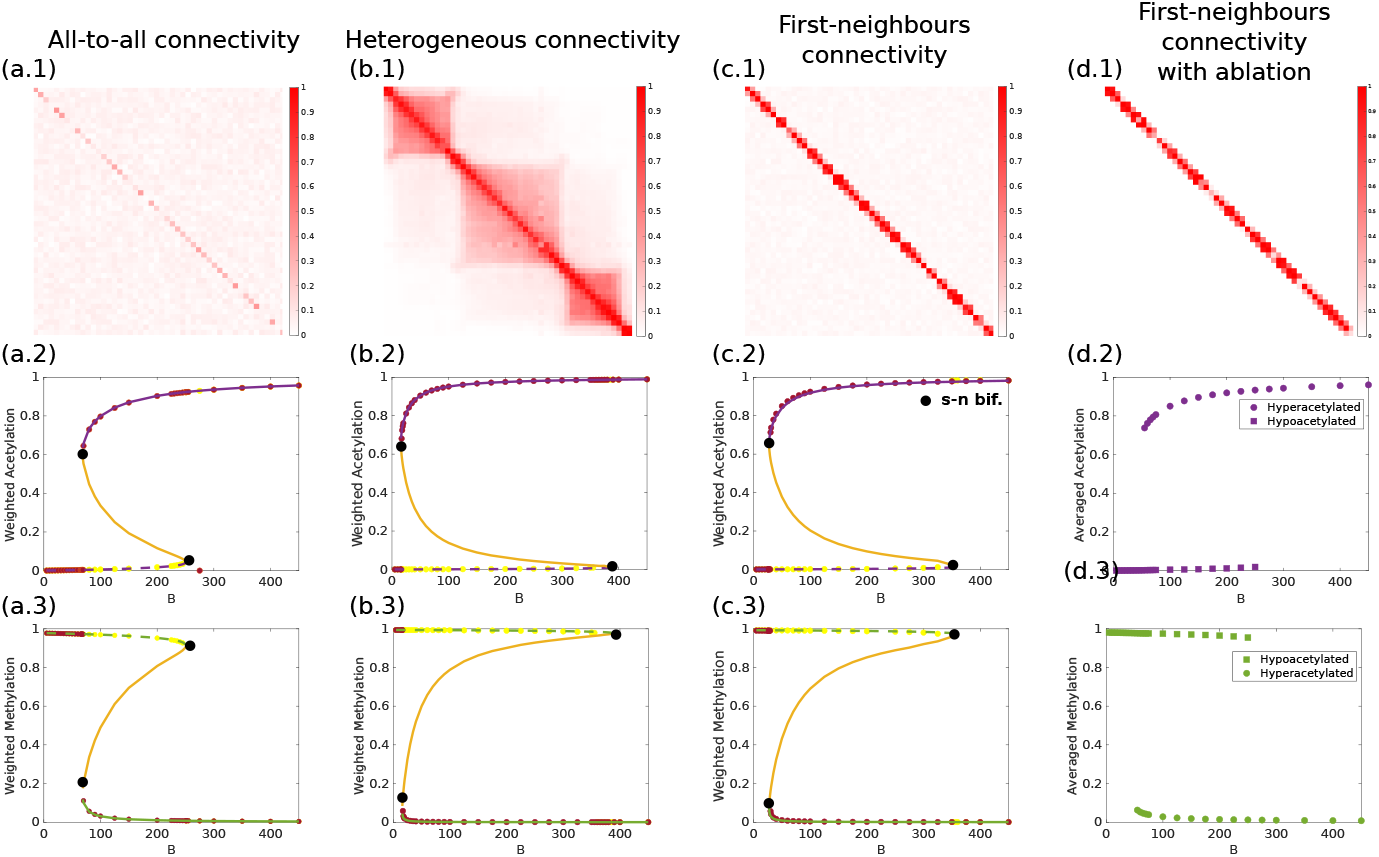
The CG method copes with different connectivity structures. We show results illustrating the robustness of our method to changes in connectivity structures. To carry out this analysis, we have used three different connectivities, namely, an all-to-all connectivity structure (a); a heterogeneous connectivity (b), as the ones detected in Hi-C measurements (see Fig. 2 in [21]); and a first-neighbors topology (c), including a case of first-neighbors with ablation (d). For simplicity, we have focused on the analysis of the mean-field limit system. Specifically, we have performed a bifurcation analysis with the parameter *B* as the control parameter. In all three cases, we show that the steady-state of the collective variables (weighted methylation and weighted acetylation) exhibits an excellent agreement between the microscopic dynamics (red and yellow dots) and the predictions of the coarse-grained system. Specifically, for different values of the DNA damage, *B*, we have let the microscopic system reach its steady state,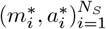. We have then plotted the steady-state value of the average methylation and average acetylation, given by 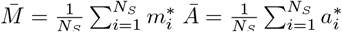. Legend: dashed lines correspond to hypoacetylated landscapes and solid lines, to hyperacetylated landscapes. The black dots represent the occurrence of a saddle-node (s-n) bifurcation.

Furthermore, as we observe comparing Figs. 4(a), (b), and (c), the connectivity pattern associated with longer-range interactions corresponds to the narrowest bistability region. By contrast, the other two patterns, heterogeneous and first-neighbors exhibit wider bistability regions. Our results imply that the width of this region does not directly correlate with the robustness of bistability. However, our robustness analysis (see Fig. 3) proves that the parameter 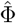, associated with the matrix W, influences robustness. Table 1 shows that 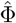 is a predictor of the range in which bistability exists.

**Table 1.** 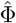 **values**. We show that the value of 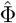 for the specific matrices used in Figs. 4(a), (b), and (c).

| Connectivity patterns | $\hat{\Phi}$ |
| --- | --- |
| All-to-all | 4.7845 |
| Heterogeneous | 72.2203 |
| First-neighbors | 38.4737 |

Finally, our model also predicts that when the intensity of the connections between bins is either too strong or too weak, the ability of our system to exhibit bistable behavior is compromised (see Supplementary Materials Fig. S4)

### Exploring epigenetic tipping points under realistic heterogeneous connectivity patterns

We have considered a pattern of connectivity that reproduces the features observed in Hi-C interaction maps. At the scale of topologically associating domains, the interaction map is heterogeneous [21, 31]. It exhibits sub-regions that are tightly connected, while the connections between such sub-regions are much weaker or infrequent. It can thus be considered as an intermediate case between the first-neighbors and the all-to-all cases: the overall level of connectivity is much higher than in the first-neighbors pattern but not as dense as in the all-to-all connectivity. To analyze this case in the context of our CG methodology, we have generated a matrix (see Fig. 4(b)) with the features of that obtained from Hi-C data (see Fig. 2 in [21]). Using this heterogeneous connectivity pattern, we have derived the corresponding CG system and performed the bifurcation analysis to show that this connectivity pattern also supports tipping points induced by competition for HDAC. The results are shown in Fig. 4(b). As in the case of the other connectivity patterns (Figs. 4(a) and (c)), the CG approximation accurately predicts the critical behavior of the microscopic system. We conclude, therefore, that the CG method can be accurately applied to realistic heterogeneous connectivity patterns.

### Bistable chromatin without long-range interactions

We have further pursued an *ablation study*, where we have used a pure, ablated first-neighbor (AFN) interaction pattern, i.e., one where *w*_*ij*_ strictly equal to zero for all non-first-neighbor interactions (see Supplementary Materials, Section II). Since W now has entries *w*_*ij*_ = 0, the CG method cannot be applied, as now it is not granted that W has a dominant eigenvector with all its components strictly positive (see Supplementary Materials). However, we can still solve numerically the microscopic dynamics, Eqs. (1)–(2). The results are shown in Fig. 4(d). Our simulations show that the system with the AFN matrix still exhibits bistability, although the bistability region becomes narrower when compared to the non-AFN first-neighbor chromatin architecture (cf. Fig. 4 (c.2) and (d.2)), suggesting that the latter’s background of low-intensity, long-range interactions stabilizes bistable chromatin. This is an interesting result regarding bistable chromatin. Most of the current models in the literature require long-range interactions as a necessary condition for bistability. The analysis with AFN interactions shows that our model does not require all-to-all or long-range interactions to support bistable chromatin. Since our model bins the genomic region of regions five nucleosomes in width (approximately), first-neighbor interactions in our model mean connections in the range of ~750 bases. Our model requires no-longer range interactions for bistability.

### Epigenetic early warning signals (EWS)

The sensitivity analysis provides a heterogeneous landscape of sensitivity, where some parameters (or combinations thereafter) are shown to affect deeply the robustness of the hyper- and hypo-acetylated states, whereas others barely have an effect (see Fig. 3).

This analysis allows us to identify which parameters can synergise positively (i.e., facilitating the tipping point) or negatively (i.e., hindering the tipping point) with the competition for HDAC modifiers. Hereafter, we illustrate how our sensitivity analysis identifies candidates to be EWS i.e., observable properties of the system that provide a warning of an impending abrupt transition in the system.

As indicated in Fig. 3, we will focus our analysis on two of the factors that affect the kinetics of HMs and HACs, *k*_1_*e*_2_ and *k*_7_*h*_0_, respectively. These quantities regulate the rate at which HMs and HACs form complexes with their corresponding substrates.

These processes depend on the availability of cofactors that act as donors of the methyl and acetyl groups. The abundance of such cofactors, S-Adenosyl methionine (SAM) and Acetyl-coA (AcoA), respectively, modulate the rate of the corresponding chromatin modifier, and it therefore provides a natural (and biologically relevant) way of changing these parameters. To tackle this, we have modified the microscopic model and derived the corresponding CG model to account for the effects of the cofactors. The details of the derivation are given in the Supplementary Materials.

By inspection, the CG equations for the model with HM- and HAC-cofactor regulated can be obtained from Eqs. (5)–(6) by making the substitutions *k*_1_ → *k*_1_*ŝ*_*c*_ and *k*_7_ → *k*_7_*â*_*c*_, where the *effective cofactors â*_*c*_ and *ŝ*_*c*_ are

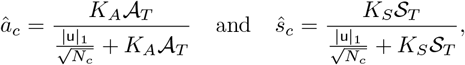

respectively. Constants *K*_*A*_ and *K*_*S*_ are the affinities and 𝒜_*T*_ and 𝒮_*T*_ are the levels of AcoA and SAM, respectively. The CG model shown above (see *Coarse-grained equations*) is the limit of the model modulated by abundance cofactor when the cofactor level tends to infinity (see Supplementary Materials). The results of our analysis varying the effective cofactors are shown in Fig. 5.

**Fig 5.**
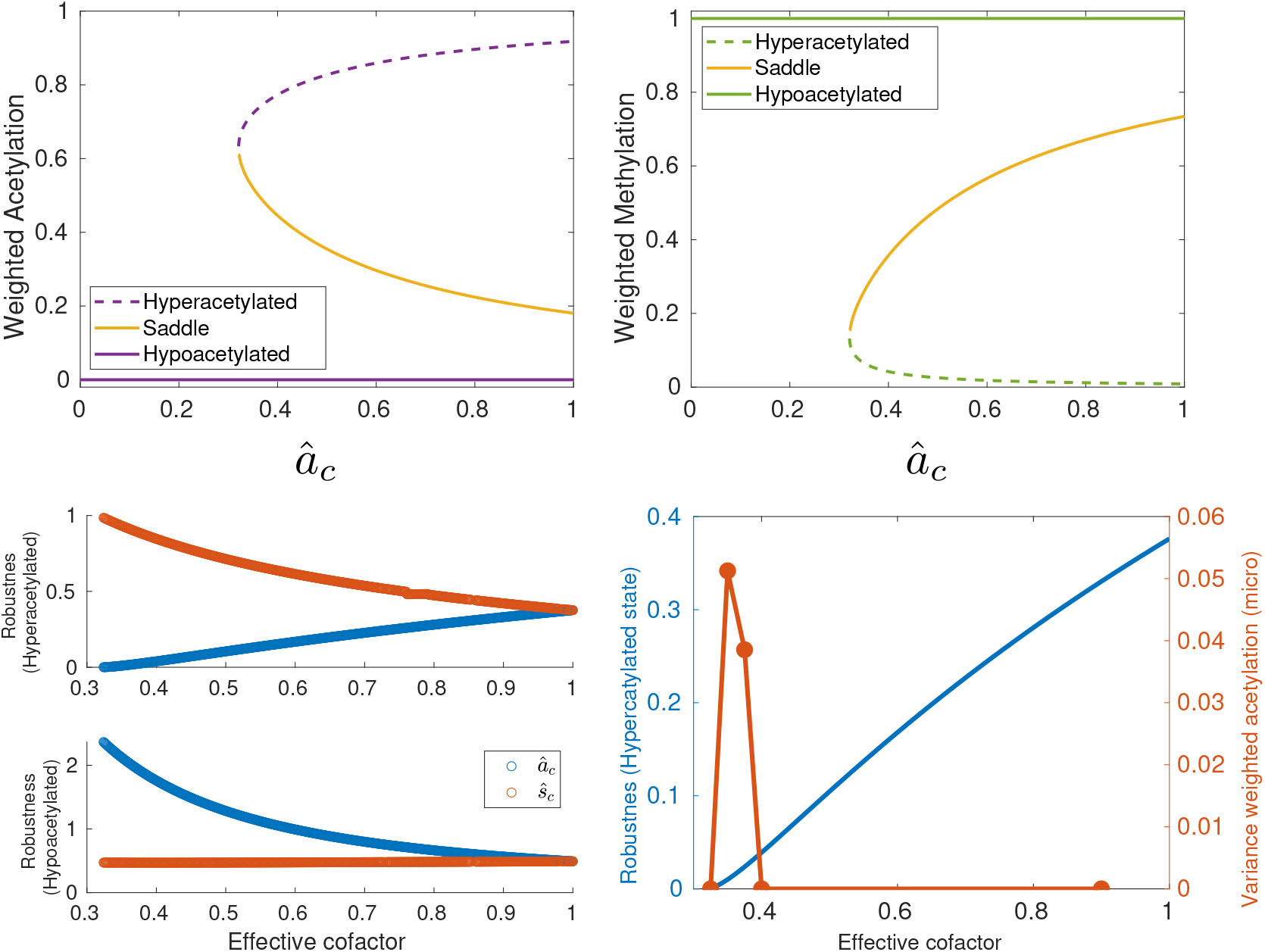
Cofactors of chromatin modifiers as early warning indicators of tipping points in epigenetic landscapes. The sensitivity analysis of the robustness of the stable ELs (see Fig. 3) indicates that the changes in the levels of metabolic cofactors SAM and AcoA (associated with the HMT and HDAC modifiers, respectively) alter the robustness of the stable ELs. Here we show that the mean-field limit of the CG approximation predicts a tipping point as AcoA availability decreases. The analysis of the stochastic CG approximation shows how the variation of these cofactors affects the robustness of the hyper- and hypo-acetylated ELs. Finally, we show that the predictions of the CG approximation accurately reproduce the behavior of the microscopic benchmark. This analysis has been done with the all-to-all connectivity.

Studying first the behavior of the CG system, as predicted by our sensitivity analysis, the variation of the effective cofactors affects the robustness of the hyper- and hypo-acetylated states. Focusing on the mean-field limit of the CG dynamics, Fig. 5 shows that reduction of the AcoA effective cofactor, *â*_*c*_, induces a saddle-node bifurcation that catastrophically destroys the hyperacetylated state. Results regarding the stochastic system are consistent with the deterministic ones. Figure 5 also shows that the robustness associated with the hyperacetylated state decreases until vanishingly small values are reached, heralding the tipping point where the hyperacetylated state disappears. It is worth noting that the critical value of the stochastic tipping point is equal to the mean-field bifurcation critical value of *â*_*c*_. By contrast, as predicted by our sensitivity analysis (see Fig. 3), reduction of the SAM effective cofactor increases the robustness of the hyperacetylated state.

More importantly, our CG analysis predicts the critical behavior of the full microscopic stochastic system. Figure 5 shows simulation results for the microscopic system as the AcoA effective cofactor decreases. Specifically, we have plotted the variance over several sample paths. We can see that as the value of the cofactor approaches the critical value at which the tipping point occurs, i.e., where the robustness of the hyperacetylated state becomes vanishingly small, the variance experiences a very pronounced peak. This behavior (vanishingly small barriers and a peak in the variance) is exactly what one expects at a tipping point [8]. Beyond that, loss of AcoA interacts synergistically with competition for HACs to induce a tipping point in the system. Since the levels of AcoA are (experimentally) observable, they are candidates to act as EWS for epigenetic tipping points.

## Discussion

Epigenetic landscapes (EL) stability is one of the most important factors in maintaining gene expression patterns in differentiated cells, thereby preserving their specific identities and functions. Recently, competition for chromatin modifiers between different epigenomic substrates (e.g., DSBs vs. histone modification-regulated gene expression) has been proposed as a key factor that can destabilize and cause loss of epigenetic information from EL [2, 5, 6]. Here, we developed a first-in-class general framework to analyze tipping points in large ELs. By measuring the sensitivity and robustness of such epigenetic tipping points, we reveal that the availability of metabolic cofactors can operate as a *bona fide* EWS able to anticipate critical transitions in ELs.

We first formulated a theoretical framework and applied it to a relatively simplified epigenetic regulatory system, where a dynamic change between trimethylation and acetylation at H3K27 underlies the switching between repressive and active genomic regions [18]. To analyze the stability of large-scale EL patterns, our framework includes a pipeline where a coarse-grained (CG) description of the system in terms of a low-dimensional system (two-dimensional, in the case of the system studied here) collective observables was obtained using the spectral reduction dimension [32, 33]. Subsequently, bifurcation theory has been used to study the CG system in the mean-field limit [Fig. 6(a)] while asymptotic techniques, in particular the WKB approximation, were used to study the corresponding Fokker-Plank equation in the stochastic diffusion limit. This analysis enabled the definition of robustness measures for stable ELs in the multi-stable regime [Fig. 6(b)]. The prediction that the large-scale transitions in ELs associated with chromatin modifier competition occur through tipping points was consistent in both the deterministic and stochastic analyses. These predictions were tested against the benchmark of the microscopic model and were found to accurately reproduce the behavior of the model [see Fig. 6(c)-(h)].

**Fig 6.**
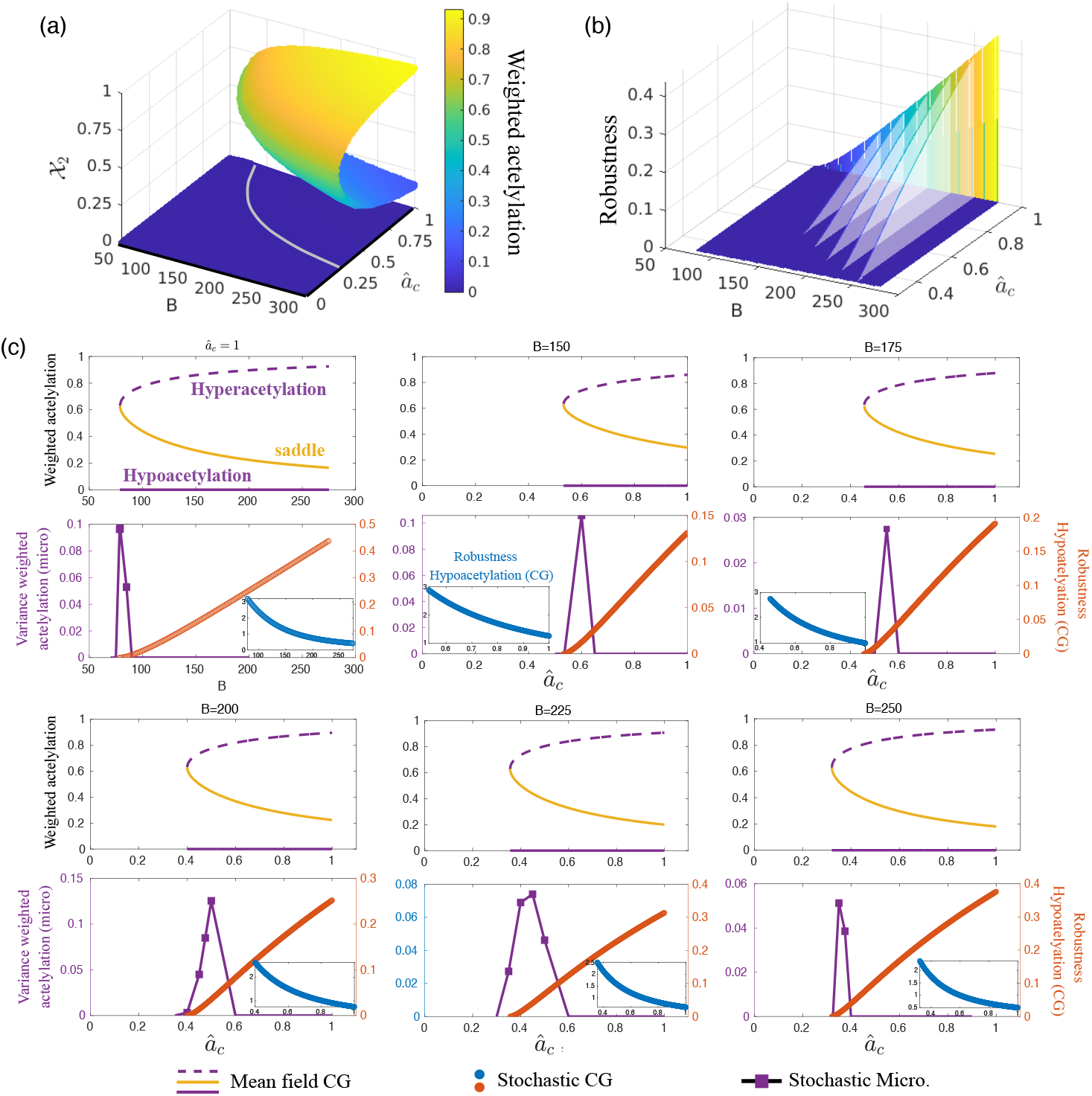
The coarse-grained (CG) approximation accurately predicts the critical behavior of the microscopic system. (a-b) CG analysis (mean-field limit and stochastic diffusion limit, respectively). These analyses in terms of the collective observables (specifically, weighted acetylation) show that competition for chromatin modifiers (HDACs in this specific case) induces a tipping point behavior (c). The analysis of CG stochastic model allows us to characterise the robustness of the stable ELs. When approaching the tipping point, the robustness of the EL becomes vanishingly small. The microscopic system exhibits a tipping point at a critical value of the control parameter that is accurately predicted by the CG approximation, as shown in (c), where the variance of the microscopic system (see purple lines) is observed to acutely increase in the vicinity of the critical point predicted by the CG approximation (orange circles). The sensitivity analysis of the CG robustness predicts that reduced levels of AcoA can facilitate the tipping point, as lowering AcoA levels decreases the robustness of the EL (b). This prediction is also faithfully fulfilled by the microscopic system, as shown in (d)-(h). This analysis has been performed with the all-to-all connectivity matrix.

By performing a parameter sensitivity analysis for the robustness metric, we were able to extract more information from the analysis of the stochastic CG approximation. This analysis informed us about the parameters whose variation has a stronger impact on the robustness of a stable EL. In other words, we can determine which parameters are more likely to facilitate the occurrence of a tipping point. Conversely, we can also determine which parameters increase the robustness of the EL and thus prevent the occurrence of the tipping point (see Fig. 3). Importantly, the robustness sensitivity analysis enables us to articulate a biologically feasible mechanism involving SAM and acetyl-CoA – the universal donors that provide methyl and acetyl units for histone modification – to control the robustness of specific stable ELs within the bistable regime [3, 23]. The predictions of the CG approximation again closely and accurately matched the behavior of the microscopic benchmark when we modified the microscopic model to explicitly account for the levels of SAM and acetyl-CoA and derived the corresponding CG approximation (Figs. 5 and 6).

Our framework, which predicts that acetyl-CoA and SAM are key regulators of the robustness of specific ELs, supports the hypothesis that the levels of certain metabo-epigenetic cofactors can be used as EWS to anticipate epigenetic tipping points driven by competition for chromatin modifiers. For example, since reduced levels of acetyl-CoA indicate reduced robustness of the hyperacetylated EL, the status of acetyl-CoA may help to predict the tipping point for initiation of an EL transition (that is, the lower the acetyl-CoA levels, the closer the tipping point). The abundance and/or distribution of metabolo-epigenetic cofactors changes significantly in response to several physiological or pathological conditions in extracellular and subcellular reservoirs [44–50]. Acetyl-CoA levels rise and fall with nutrient abundance, oncogenic signaling, and microenvironmental conditions, and such changes in acetyl-CoA abundance can regulate histone acetylation at specific loci and the expression of distinct sets of genes. Acetyl-CoA status can also be coordinated also with chromatin structure and function through spatiotemporal control of nuclear acetyl-CoA concentration via post-translational modification and/or dynamic chromatin recruitment and localization of acetyl-CoA-producing enzymes [44–50]. Serine starvation, a common microenvironmental stress in breast cancer that leads to a marked and rapid insufficiency of the nucleocytosolic reservoir of acetyl-CoA, has recently been shown to deplete H3K27 acetylation and induce a global transition to an hypoacetylated EL [51]. Given that the acute depletion of acetyl-CoA culminates in epigenetic reprogramming and silencing of a plethora of genes important for defining cell lineage commitment, such as those involved in the cellular response to estrogen [51], these results experimentally support our proposal that, in response to small changes in the environmental availability of metabolic cofactors (e.g., acetyl-CoA) near a tipping point, critical transitions may occur driven by competition for chromatin modifiers.

Accordingly, the provision of acetate –an alternate source of acetyl-CoA– is sufficient to restore EL and prevent estrogen receptor (ER) silencing under serine-free conditions [51]. In analogy to acetyl-CoA, systemic and local changes in SAM-producing pathways may also contribute to the variability of histone methylation patterns and chromatin regulation of different functional cellular states in tissues and organs. The dietary and extracellular availability of methionine and serine significantly affect SAM levels and histone methylation, and we are beginning to realize that nuclear SAM production may also be important for the function of specific chromatin-associated complexes that regulate histone methylation [44–50]. The growing evidence that acetyl-CoA and SAM are key messengers that translate nutrient-dependent and compartment-specific metabolic signals into chromatin modifications not only provides a robust biological basis for our proposal of metabolic cofactors as EWS that can predict the onset of EL global transitions but also suggests metabolic strategies (e.g., dietary interventions, targeted modification of metabolic pathways) to avoid them.

Our sensitivity analysis shows that in addition to metabolic (co)factors affecting the activity of chromatin modifiers, the structure and intensity of the interactions are also factors that quantifiably affect the robustness of bistable EL (summarized within the parameter 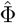, as shown in Fig. 3). To address this issue in more detail, we considered three connectivity patterns, namely all-to-all, heterogeneous, and first-neighbours considering also a particular case with ablation. In addition to demonstrating that our CG method can handle a wide variety of interaction patterns and accurately predict the critical behavior of the system (Eqs. (1)–(2)), an ablation study shows that our model does not require long-range interactions to generate bistable chromatin. Interactions in the range of~ 5 nucleosomes (~ 750 base pairs) are sufficient to confer such a property. Our model uniquely approaches the activity of chromatin modifiers using enzyme kinetics rather than (recruited, nonlinear) binding/unbinding kinetics [19, 23, 29, 30].

The nonlinearity that this feature introduces into the kinetics of chromatin modifiers (see Eqs. (1)–(2) and Supplementary Materials), facilitates the robust production of bistable behavior. In line with the results of previous models [19, 23, 29, 30], the introduction of longer-range interactions (even weak ones) helps to increase the robustness of bistable chromatin.

Finally, there are several ways to extend our current framework to account for tipping points in multistable ELs. One obvious extension would be to consider more chromatin modifiers that generate more complex EL, particularly those that involve rugged, non-uniform patterns of chromatin modifications [5] This would allow computational testing of recently proposed models of epigenetic information storage in which histone modifications such as H3K27me3 become dysregulated over time [5, 52, 53].

As EL transitions are beginning to emerge as driving mechanisms for critical transitions in epithelial-hybrid-mesenchymal cell fate determination that accompany metastatic, drug-resistance phenomena during cancer progression [54, 55]) and disruption of developmental genes during aging [5, 52, 53, 56], our mathematical methodology opens a new avenue for obtaining improved EWS of more complex epigenetic tipping points for both theoretical and practical purposes. Importantly, the ability to anticipate critical EL transitions using the abundance of certain metabolic cofactors as EWS may provide predictive biomarkers and uncover new opportunities for therapeutic restriction of pathological cell fate decisions, but also advance the application of therapies to restore cellular identity in post-injury regeneration. In this regard, exogenous supplementation of a few metabolites, including methionine and SAM, that mimic an endogenous one-carbon metabolomic wave that occurs during early cell transitions in differentiation has been shown to increase global acetylation marks and prime cell identity changes to promote faster regeneration after muscle injury in young and aged mice [57]. These findings strongly suggest that our current proposal of epigenetic metabolites as EWS for critical EL transitions may provide a translatable strategy to metabolically redirect cell identity and restore cell and tissue function during disease and aging.

## Supporting information

### S1 Appendix. Supplementary materials

This appendix contains full details regarding the model formulation and detailed derivations of the coarse-grained model reduction.

## Acknowledgments

T.A. is grateful to the Isaac Newton Institute for Mathematical Sciences for support and hospitality during the program *Mathematics of movement: an interdisciplinary approach to mutual challenges in animal ecology and cell biology* during the final stages of this work under the EPSRC grant number EP/R014604/1. This work has been funded by the State Research Agency (AEI), through the Severo Ochoa and Maria de Maeztu Program for Centers and Units of Excellence in R&D (CEX2020-001084-M). We thank CERCA Program/Generalitat de Catalunya for institutional support. J.S. and T.A. have been funded by grant PID2021-127896OB-I00 funded by MCIN/AEI/10.13039/501100011033 ‘ERDF A way of making Europe’. The work in the Menendez laboratory is supported by the Ministerio de Ciencia e Innovaciòn (MCIN, grants PID2019-104055GB-I00 and PID2022-141955OB-I00 to J.A.M., Plan Nacional de I+D+i, funded by the European Regional Development Fund, Spain) and the Emerging Research Group SGR 2021 01507 from the Agència de Gestiò d’Ajuts Universitaris i de Recerca (AGAUR, Generalitat de Catalunya).

